# Isolation-By-Distance-and-Time in a Stepping-Stone model

**DOI:** 10.1101/024133

**Authors:** Nicolas Duforet-Frebourg, Montgomery Slatkin

**Affiliations:** Department of Integrative Biology, University of California Berkeley, Berkeley CA 94720

## Abstract

With the great advances in ancient DNA extraction, population genetics data are now made of geographically separated individuals from both present and ancient times. However, population genetics theory about the joint effect of space and time has not been thoroughly studied. Based on the classical stepping–stone model, we develop the theory of Isolation by Distance and Time. We derive the correlation of allele frequencies between demes in the case where ancient samples are present in the data, and investigate the impact of edge effects with forward–in–time simulations. We also derive results about coalescent times in circular/toroidal models. As one of the most common way to investigate population structure is to apply principal component analysis, we evaluate the impact of this theory on plots of principal components. Our results demonstrate that time between samples is a non-negligible factor that requires new attention in population genetics.

## 1 Introduction

Geography plays a central role in the pattern of genetic differentiation within a species. Seminal work on describing the evolution of continuous populations was done by Wright and Malécot. They studied genetic differentiation and inbreeding in continuously distributed population (Wright, 1943; Malécot 1948). The resulting idea is that, under the assumption of local dispersion, genetic differentiation accumulates with distance. This pattern of genetic structure is called Isolation–By–Distance (IBD), which is detected by computing measures of differentiation such as *F_ST_* (Wright, 1943; Nei, 1973; Weir and Cockerham, 1984), or correlation coefficients (Malécot 1955; Kimura and Weiss, 1964). Understanding the effect of geographic distance on population structure is an important task for population geneticist, as it is a source of neutral genetic variation (Slatkin, 1985; Rousset, 1997). Furthermore, IBD has been observed in humans and many other species (Sharbel et al., 2000; Castric and Bernatchez, 2003; Ramachandran et al., 2005; Hellberg, 2009; Karakachoff et al., 2015).

The role of geography in neutral genetic variation has been widely studied partly because of the existence of many population genetic studies of individuals living at the present time and sampled in different locations. Because of the development of methods for sequencing DNA from fossils, genomes of individuals alive at previous times are now available to bring new information about the evolutionary processes that affected a species in the past. Since the first studies of ancient DNA (aDNA) three decades ago (Higuchi et al., 1984; Pääbo, 1985), techniques to retrieve DNA molecules from ancient bones have tremendously developed (pääbo, et al., 2004).

In modern evolutionary biology, the similarity of differentiation in space and time is acknowledged (Depaulis et al., 2009; Andrello et al., 2011; Teacher et al., 2011). Theoretical developments predict the effect of time on *F_ST_* and related quantities (Skoglund et al., 2014). Epperson (2000) studied patterns of isolation by distance and time in ecology by using stochastic spatial time series and Identity by descent probabilities However such theoretical studies remain scarce.

The effect of separation in time can be studied using classical statistical methods in population genetics, such as principal component analysis (PCA) (Patterson et al., 2006). PCA is widely used to determine relatedness between individuals, and is a convenient way to represent geographic patterns (Novembre et al., 2008). But PCA can also capture the differentiation between ancient and modern samples: the percentage of variance explained by time can be expressed on the same scale as the percentage of variance explained by geography (Skoglund et al., 2014). Unfortunately, PCA does not give a complete picture of how differentiation quantities such as *F_st_* and correlations of allele frequencies evolve in time and space.

In this article we generalize the theory of IBD to allow for difference in the times at which different individuals are sampled. We call this the theory of isolation by distance and time (IBDT). We base our work on the stepping–stone model of Kimura (1953) and add to the theoretical results known for this model (Kimura and Weiss, 1964; Weiss and Kimura, 1965; Maruyama, 1971a; Nagylaki, 1983; Cox et al., 2002; De and Durrett, 2007). We start by briefly reviewing the original results for the infinite stepping– stone model at equilibrium and the decay of correlation of allele frequencies with distance. Then, we extend the original work to derive the correlation between individuals separated by distance and time. We perform simulations that show the validity of the analytic results, even in the case of a finite number of populations where some demes are subject to edge effect. We also derive the expected coalescence times between samples separated by time and space in circular and toroidal models (Slatkin, 1991, 1993). Finally we consider the consequences of IBDT on PCA in the common case of a dataset made of a large proportion genomes from present–day individuals and few ancient genomes.

## 2 The stepping–stone model

The stepping–stone model describes the distribution of allele frequencies in an infinite set of demes in different locations of the space represented by Cartesian coordinates. We start by describing the 1-Dimensional case. Let *p*(*k*) be the frequency of one allele at a bi-allelic locus in population *k* and 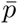 be the overall allele frequency. In each generation, *p*(*k*) is updated with the following three steps (Crow et al., 1970):

- Exchange a proportion *m_i_* of migrants with demes at a distance *i*.
- Exchange a proportion *m*_*∞*_ of migrants with a deme that has fixed allele frequency 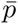. The meaning of this step is discussed later.
- Sample gametes of the next generation in the population.

In the case considered by Kimura and Weiss (1964), migrants are exchanged only between neighboring locations in the first step, so that *m_i_* = 0, *i* > 1. The second step consists of exchanges of migrants with an external population at rate *m*_*∞*_. This event is equivalent to reversible mutations occurring. The formulation of the model states that every locus is bi-allelic, and the number of loci is fixed. As a consequence, the mutation model is a reversible mutation model with probability *m*_*∞*_, and *m_i_* > > *m*_*∞*_. Random sampling of step 3 is represented by a random change in the allele frequency *ϵ*(*k*), with *E*[*ϵ*(*k*)] = 0, and *E*[*ϵ*(*k*)^2^] = *p*(*k*)(1 - *p*(*k*))/2*N_e_*, where *N_e_* is the effective population size of a deme (Wright, 1940; Kimura and Crow, 1963).

Our interest is in the changes in allele frequency in one generation. We consider 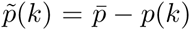, the deviation from the average frequency. Given these three steps,

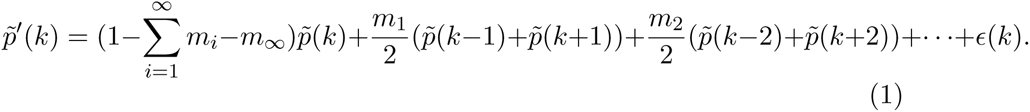

To simplify the notation, we define the operators *S* and *L*,

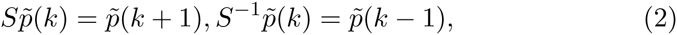

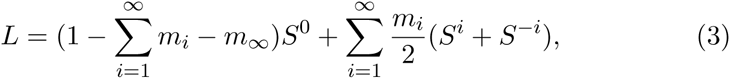

so that,

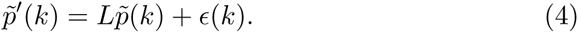

The quantity of interest in this model is the correlation of allele frequencies between two demes at locations *k*_1_ and *k*_2_. Let *r*(*k*) be the correlation coefficient of allele frequencies between populations that are *k* steps apart. Assuming equilibrium, we have

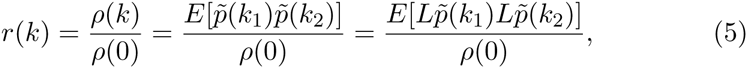

where *ρ*(*k*) is the covariance in frequencies in demes *k* steps apart. The mathematical treatment of equation (5) by Weiss and Kimura (1965) using the spectral representation of a correlation (Doob, 1953) gives the general formula

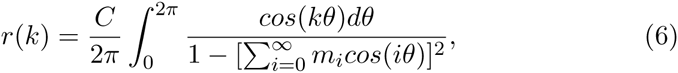

where *C* is the normalizing constant.

Equation (6) can be approximated by an exponential function of *k*:

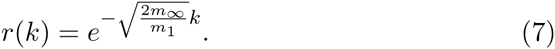

This simple formula conveys the important idea that in one dimension, the correlation of allele frequencies between populations decays exponentially with distance. In the 2–Dimensional and 3–Dimensional cases, the correlation function is more difficult to approximate. Using modified Bessel function, it is shown that correlation at a given distance is lower in these cases than in the 1–Dimensional case (Weiss and Kimura, 1965).

## 3 Isolation–by–Distance–and–Time

### 3.1 1–Dimensional case

We are here interested in the case where genetic samples are collected from demes that are in different locations and at different times (measured in generations). Let *ρ*(*k, t*) be the covariance between allele frequencies of two demes separated by *k* steps and *t* generations. We denote the coordinates of these demes by (*k*_1_, *t*_1_) and (*k*_2_, *t*_2_), and the deviations in allele frequencies 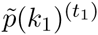 and 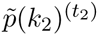 Since we assume the distribution of allele frequencies is stationary in both time (equilibrium distribution) and space (all migration rates are equal), we can consider these coordinates to be (0, 0) and (*k, t*) with no loss of generality. Following previous notation

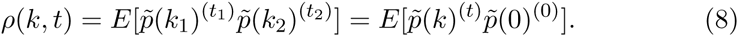

To characterize the evolution of the covariance between allele frequencies with respect to time *t*, we iteratively apply the operator *L* defined in equation (3). This operation describes the potential trajectories of an allele, and results in a quantity similar to a propagator. This process leads to

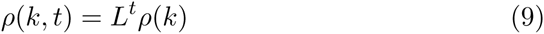

with *ρ*(*k*) = *ρ*(*k,* 0) (see Appendix A).

Let *r*(*k, t*) be the correlation between allele frequencies of two demes separated by *k* steps and *t* generations, equations (5) and (9), combined with the general formula of equation (6) gives

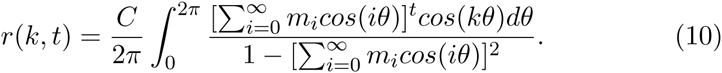

and the constant C is set such that *r*(0, 0) = 1 (Appendix B).

This equation reduces to

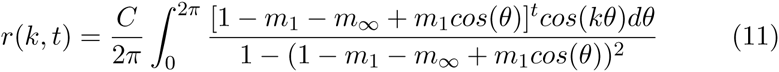

in the standard stepping–stone model, where demes only exchange migrants with their closest neighbors at rate *m*_1_/2. An exact formula for this integral can be calculated and is notable for its size and lack of utility (Appendix C).

One noteworthy feature of equation (10) is that the decay of the correlation with time is not affected by the effective population size *N_e_*. This result is different from what is expected for an isolated population: the level of differentiation as a function of the number of generations separating two samples is larger when the effective population size is small, reflecting the increased magnitude of genetic drift. However, in the particular case of an equilibrium stepping–stone model, the covariance of allele frequencies between the demes is not a function of the effective population size, a result already known in the spatial context (see equation (7)) (Kimura and Weiss, 1964). This result becomes clear when considered in terms of coalescence times. Between the time the first and second samples are taken, the trajectory of the first sample depends only on the migration process. There is no possibility of coalescence.

### 3.2 Two dimensions and more

So far, we have focused on the 1-Dimensional case for the sake of simplicity. However, it is important to investigate the decay in higher dimensions as it is common in practice to have samples taken from a 2-Dimensional or even 3-Dimensional habitat. The general formula for the correlation in higher dimensions can be obtained with no more theoretical development. In their work on the stepping–stone model, Kimura and Weiss derived a general formula for the correlation that can be extended to any number of dimensions. In their work they only gave approximations for 1, 2 or 3 dimensions as these are the practical cases. Using general formula (3.11) of Weiss and Kimura (1965), we can write the correlation 10 in 2 dimensions

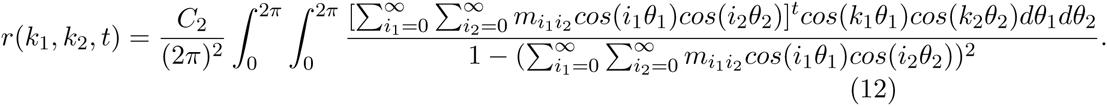

The generalization to obtain the correlation in n dimensions is straight– forward (Appendix D).

We perform a numerical integration of equation (12) to investigate the decay of correlation with distance and time in one dimension and higher. Correlation decreases as a function of distance and time in 1, 2 and 3 dimensional models (Figure 1). In addition, for equal values of the migration and mutation rates the correlation decrease is much larger with respect to time and geography in higher dimension models, consistently with previous results (Maruyama, 1970a, 1971a). Numerical integration is done using the *R* package *cubature*.

**Figure 1:**
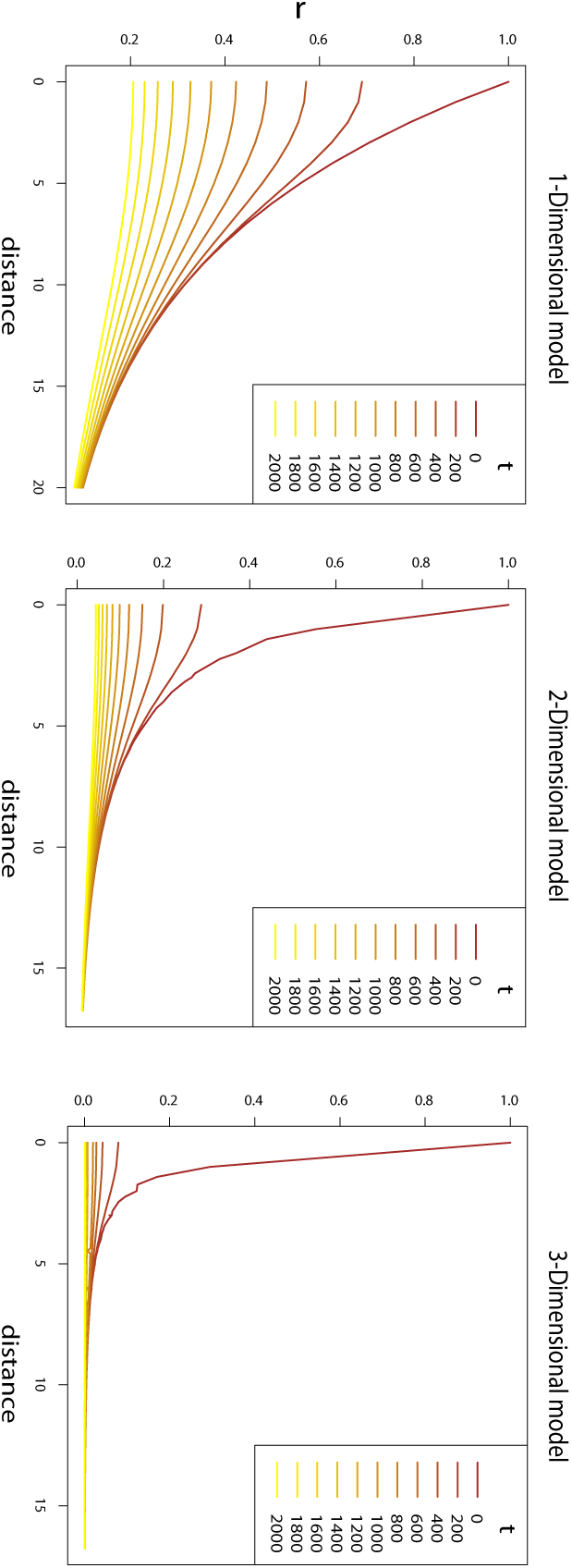
Correlation as a function of distance between demes *k* steps appart in 1, 2 and 3-Dimensional models. The correlation is evaluated for different number of generations *t* between the demes. The migration and mutation rates are used for all models, and *m*_1_ = .01 and *m*_∞_ = 4.10^−4^.

### 3.3 Simulations in one dimension and two dimensions

When considering realistic examples, a finite number of demes is present in the data. As a consequence, correlation patterns are affected by the proximity of the edge of the sampling scheme (Maruyama, 1970b). Another effect of the finite number of demes is that the overall allele frequency can drift away from the theoretical allele frequency. An alternative is to consider a finite, non-circular model, and to deal with edge issues independently (Felsenstein, 2015). To investigate to which extent the analytic theory developed in the previous section is valid in a finite stepping–stone model with temporal sampling, we perform simulations.

Backward in time simulation software such as *ms* (Hudson, 2002), or *fastsimcoal* (Excoffier and Foll, 2011), are usually used to investigate IBD in a stepping–stone models (Novembre et al., 2008). Temporal sampling can be investigated in such model by simulating gene trees where lineages from isolated demes are joined to the stepping–stone demes at a chosen time in the past (Skoglund et al., 2014). Mutations are then randomly placed on the gene tree. Such a simulation is needed to understand the influence of time and distance on genetic differentiation, but does not precisely reproduce the process of the above model which assumes reversible mutation rather than the infinite site model. The infinite site model does not have a true equilibrium for any one site, only a pseudo–equilibrium.

We wrote a C program that performs forward in time simulations. The simulation program precisely follows the model presented in the previous section. At the initial time, the allele frequencies in all the demes are equal to the allele frequencies in the outside infinite–sized population. Then the program runs for a large number of generations until the stationary distribution of the allele frequencies is reached.

In the 1-Dimensional case, we simulate 100 demes. For the 2-Dimensional case, we simulate a total of 2500 demes on a 50 × 50 grid. We assume all the demes have the same effective population size. We sample the allele frequencies at several times in the past. Correlation between demes fit very closely the theory of equations (11) and (12) provided that demes are taken sufficiently far away from the edge of the grid (Figure 2). The edge effect directly increases the correlation between demes, and is present when com paring present and ancient samples. In both 1 and 2 dimensions, the edge effect disappears in the simulations (Figure 3). As predicted by Maruyama, the edge effect is less strong with lower migration rates.

**Figure 2:**
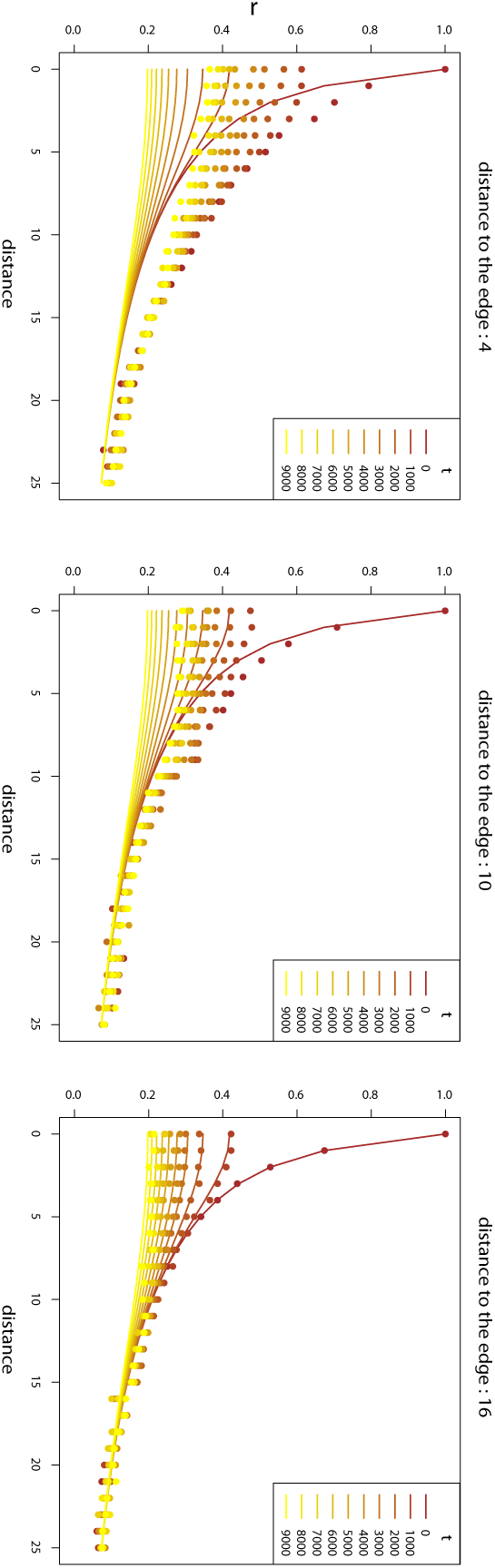
Comparison between theoretical results and simulations in the 2 dimensional case with *m*_1_ = .02 and *m*_∞_ = 10^−5^. The solid lines represent the theory prediction. The dots represent the simulation results evaluated for demes at a distance 4, 10 or 16 from the edges. Since in the simulations several pairwise comparisons between demes have the same distance in space and time, we keep the median of these pairwise correlations.

**Figure 3:**
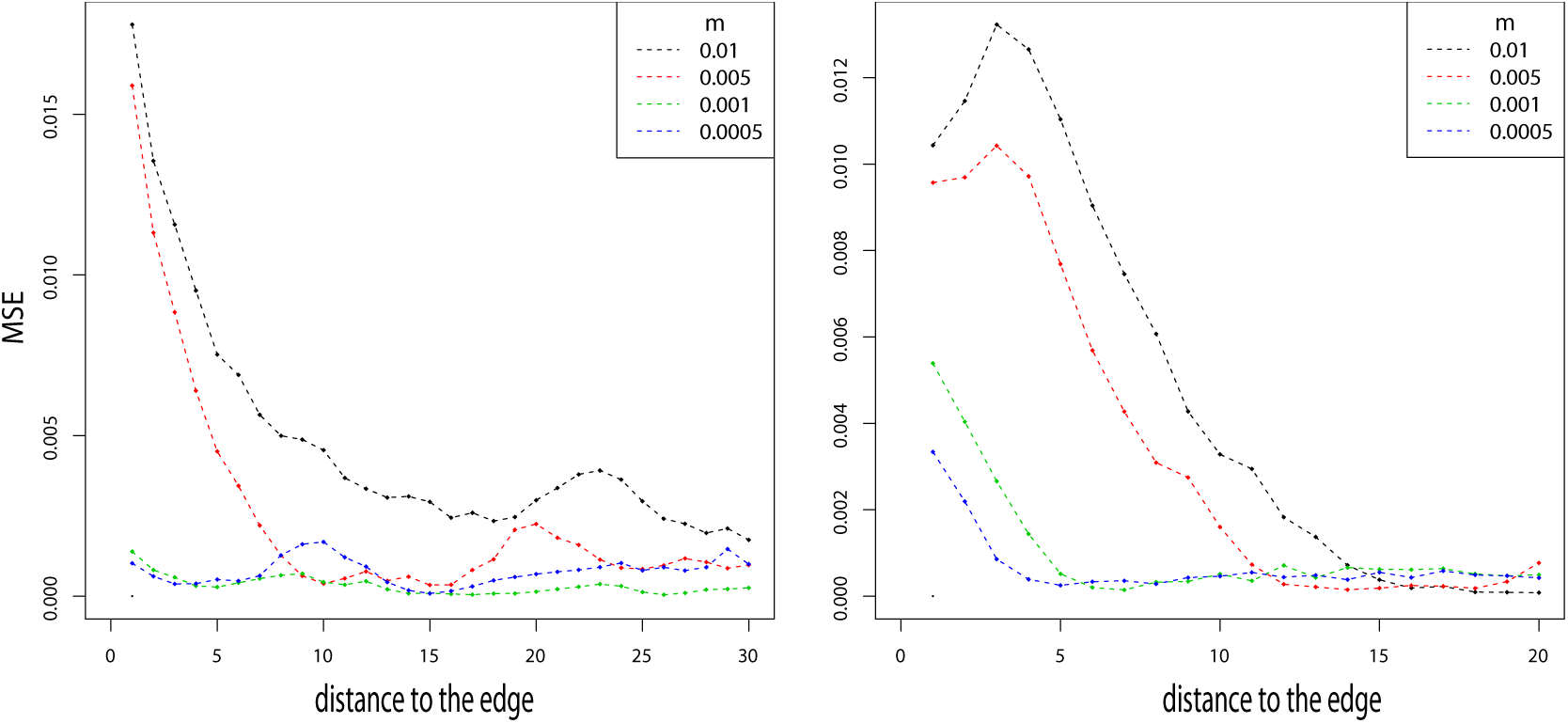
Mean squarred error between simulations and theory in 1 and 2 Dimensions as a function of the distance to the edge. The error is evaluated for *m*_∞_ = 10^−5^ and *m*_1_ = .01, .005, .001, .0005.

## 4 Coalescence times

### 4.1 Coalescence times in one dimension

Coalescence times in a stepping–stone model can be derived under some assumptions. In particular, we consider a case with migration only between neighboring demes and low mutation rate. Expected coalescence times between genes that are in different demes is a function of the locations of these demes. These coalescence times are of interest because they closely related to *F_ST_* and coefficients of identity–by–descent (Slatkin, 1991). Under the assumption of a circular 1-Dimensional stepping-stone model with *n_d_* demes, two genes *A*_1_ and *A*_2_ have an expected coalescence time

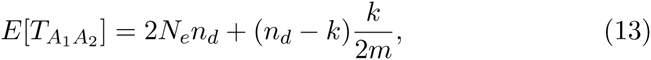

where *N_e_* is the effective population size per deme, *m* the migration rate between neighboring demes (previously *m*_1_), and *k* is the distance between the two demes (Slatkin, 1991). Considering a circular arrangement of the demes makes the analysis simpler, as only the distance between the demes matters, and there are no edge effects. In addition it has been shown that linear/planar and circular/toroidal stepping stone models are very similar when considering population away from the edges (Maruyama, 1971a,b). To study a case similar to the infinite stepping–stone model, we assume *n_d_* is large.

We extend the previous theoretical result in the case where two genes are sampled at different times. Let us assume that the sampled genes are in population *k*_1_ and *k*_2_. The number of generations between the two sampling times is *t* = *t*_1_ - *t*_2_, and we assume, with no loss of generality, that *t*_1_ = 0 and *t*_2_ = *t* generations in the past. The coalescence process between these two genes can be divided in three phases. The first phase corresponds to the genealogy that traces back to the ancestor of the present gene, called 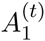, at generation *t*. This ancestor is in population 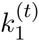. The two other parts correspond to the time until the coalescence event between 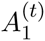 and *A*_2_. They are respectively the time until the gene 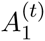 and *A*_2_ are in the same deme, then the time to the common ancestor of these two genes. This part has already been described, and the expectation is given in equation (13) (Slatkin, 1991). The expected coalescence time between *A*_1_ and *A*_2_ is then written

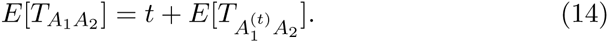

The variable 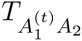 is the coalescence time between a random gene in the unknown population 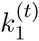 and a random gene in population *k*_2_. To represent the uncertainty about the population 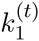, we derive the probability distribution of the position 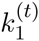 at time *t*, given position *k*_1_ at time 0. Using this probability distribution we rewrite the expectation (14) as

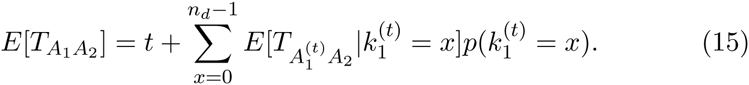

To describe the probability distribution of position 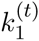 at time *t* given that a gene is in population *k*_1_ at time 0, we consider a random walk with transition matrix

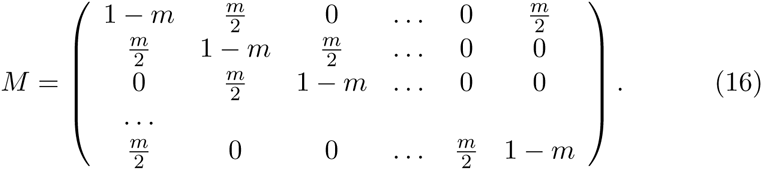

Using standard results about Markov chains (Ross et al., 1996), we know that the vector of probabilities for the position at time *t*, 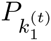 is expressed such as

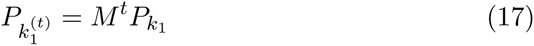

with *P*_*k*_1__ is the initial probability distribution of gene A_1_’s position. The initial probability distribution is trivial and *P*_*k*_1__ is a vector of 0 with a 1 in the 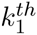 entry. Exact formula for this matrix power can be obtained using tridiagonal matrix properties (Al-Hassan, 2012). However we can also express an approximation for the probability distribution of this process at time *t*. This random process is symmetrical, centered in *k*_1_, and using classical results about Brownian motion, has a variance proportional to *t*. We can approximate the probability distribution by a Normal distribution, and

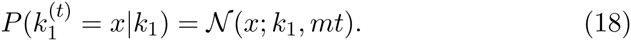

The accuracy of this approximation can be verified with simulations using equation (17). The approximation is relevant for sufficiently large values of *t*, depending on the migration rate. The expected coalescence time in a 1-Dimensional circle can then be written

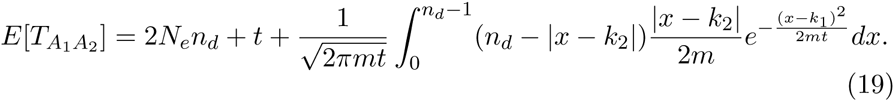

Coalescence time between genes is an increasing function of distance and time between demes (Figure 4). Asymptotically, when *t* is large, the expected time for two genes to be in the same population can be approximated by a linear function of time between the samples (Figure 4). The right part of equation (19) is the integral of a product of a positive function that depends only on the distance between demes and a Gaussian kernel with variance *mt*. As the time gets large, relatively to *m*, the Gaussian kernel becomes flat, and the integral is almost constant (Figure 4). In practice, this implies that in a population at equilibrium, the geography does not matter when the sample is very old.

**Figure 4:**
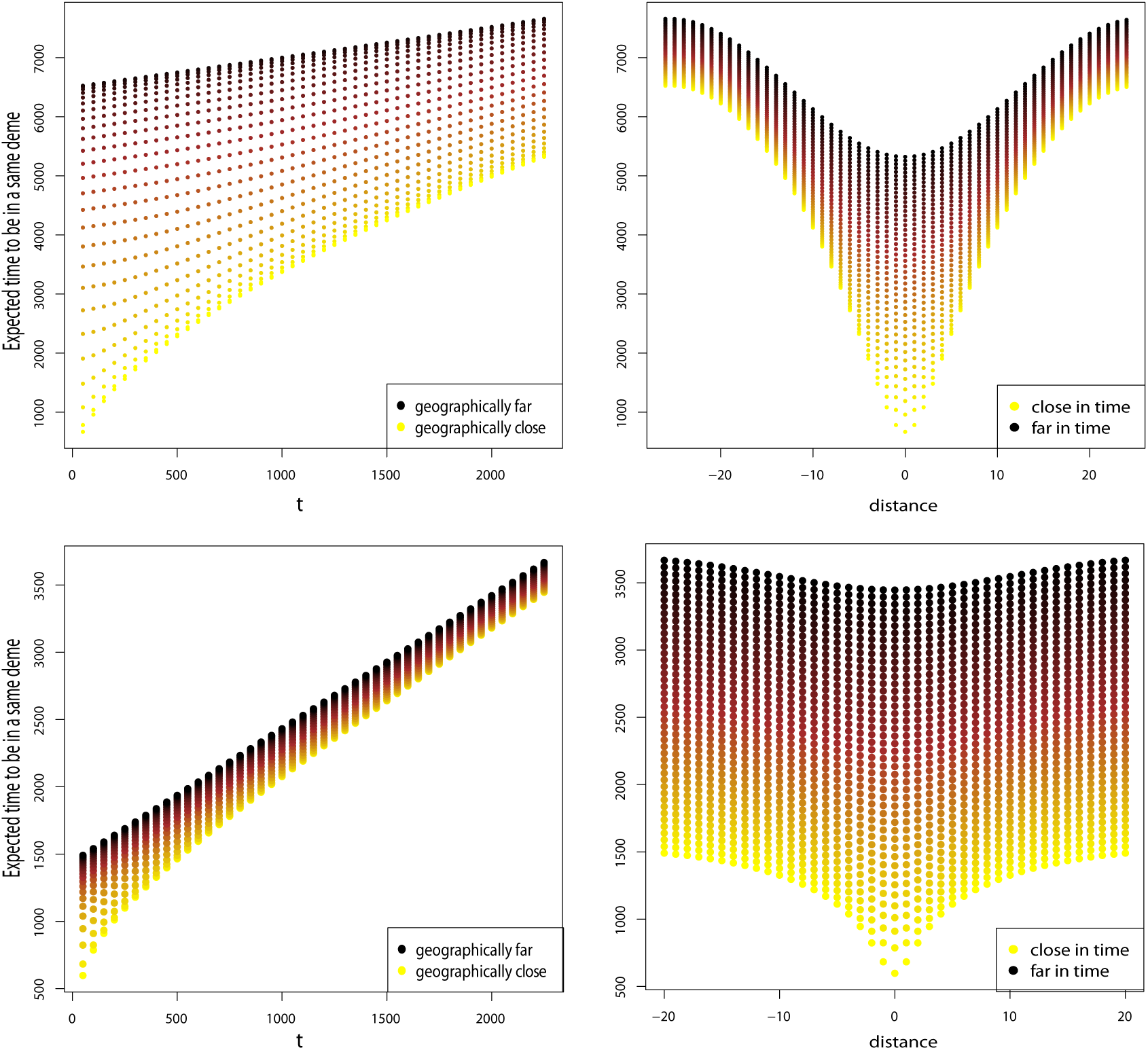
Top row: Expected time for two genes to be in a same deme in a 1–Dimensional circular stepping–stone model with *N_e_* = 100, *m* = .01, and *n_d_* = 51 demes. Bottom row: Expected time for two genes to be in a same deme in a 2–Dimensional toroidal stepping–stone model with *N_e_* = 100, *m* = .01, and *n_d_* = 51×51 demes. Left column: Expected times as a function of the time between the samples. Colors indicate the geographic distance between samples. Right column. Expected times as a function of geographic distance between the samples. Colors indicate the time between samples. Sampling consists in 45 time points evenly separated by 50 generations.

### 4.2 Coalescence times in two dimensions

In the case of a 2-Dimensional habitat with *n*_*d*1_ × *n*_*d*2_ demes, the expected coalescence time between two genes *A*_1_ and *A*_2_ is

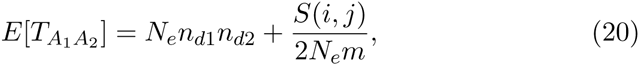

where *S*(*i, j*) is a function of *i* and *j*, the number of demes between the two genes. We assume in this case that the migration in each direction is the same.

Using the same conditioning as in equation (14), we can derive the expectation for the coalescence time of genes *A*_1_ in population *k*_1_ and *A*_*2*_ in population *k*_2_, *t* generations in the past. We have

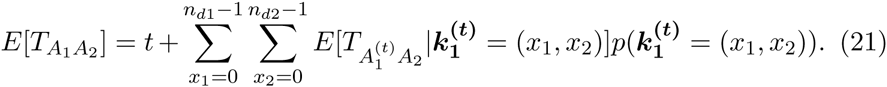

The probability distribution of the position of gene *A*_*1*_ at time *t*, 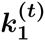 is known using the same random walk as in the 1-Dimensional case. The distribution can be approximated by a bivariate Normal distribution with mean ***k*_1_**, and covariance matrix Σ, where Σ is diagonal with terms *mt*/2 in the diagonal. In the anisotropic case where migration rate would be different in the two dimensions, *m*_1_ and *m*_2_, Σ would have *m*_1_*t* and *m*_2_*t* as diagonal terms. The evaluation of this function for samples separated in distance and time shows a similar pattern to the 1-Dimensional case (Figure 4). However for a same migration rate, the expected times for two genes to be in the same deme in the 2–Dimensional toroidal model are smaller than in the 1– Dimensional circular model. Then, if there is the same number of demes, with same effective population sizes, e.g. *n_d_N_e_* = *n*_*d*1_*n*_*d*2_*N_e_*, the expected coalescence times are smaller in the 2–Dimensional case. This result is already known when comparing samples taken at the same generation and remains true when *t* is positive (Slatkin, 1993).

## 5 Connection with PCA

Because there is a close connection between PCA and coalescence times (McVean, 2009), our results are relevant to using PCA to compare ancient and modern samples. PCA is a useful way to represent the main axes of variation in the data and has proven to be a powerful tool to infer genetic relationships when applied to ancient DNA data(Skoglund et al., 2012; Haak et al., 2015).

### 5.1 Ancient samples are shrunk towards 0

In population genetics, PCA is usually performed by computing the eigenvectors, and eigenvalues of the matrix of covariances in the genotypes of different individuals. Although there are other ways to compute principal components, this one is convenient in population genetics because the number of variables is usually larger by several orders of magnitude than the number of samples. The effect of differences in the sampling times can be evaluated using the dependence of the covariance matrix described by equation (10). To illustrate, consider a 2-Dimensional even repartition of 10 × 10 demes, and ancient samples taken in several randomly chosen demes at *t* = 1000 generations in the past (Figure 5A). By calculating the theoretical covariance matrix and its first two eigenvectors, we obtain the first two principal components that reproduce geography of the demes (Novembre et al., 2008; Engelhardt and Stephens, 2010). Figure (5B) shows that principal components mimic the geography of the present demes, but ancient demes are not superposed on the corresponding present-day sample from the same deme. Instead, ancient samples move towards the center of the first and second principal components.

**Figure 5:**
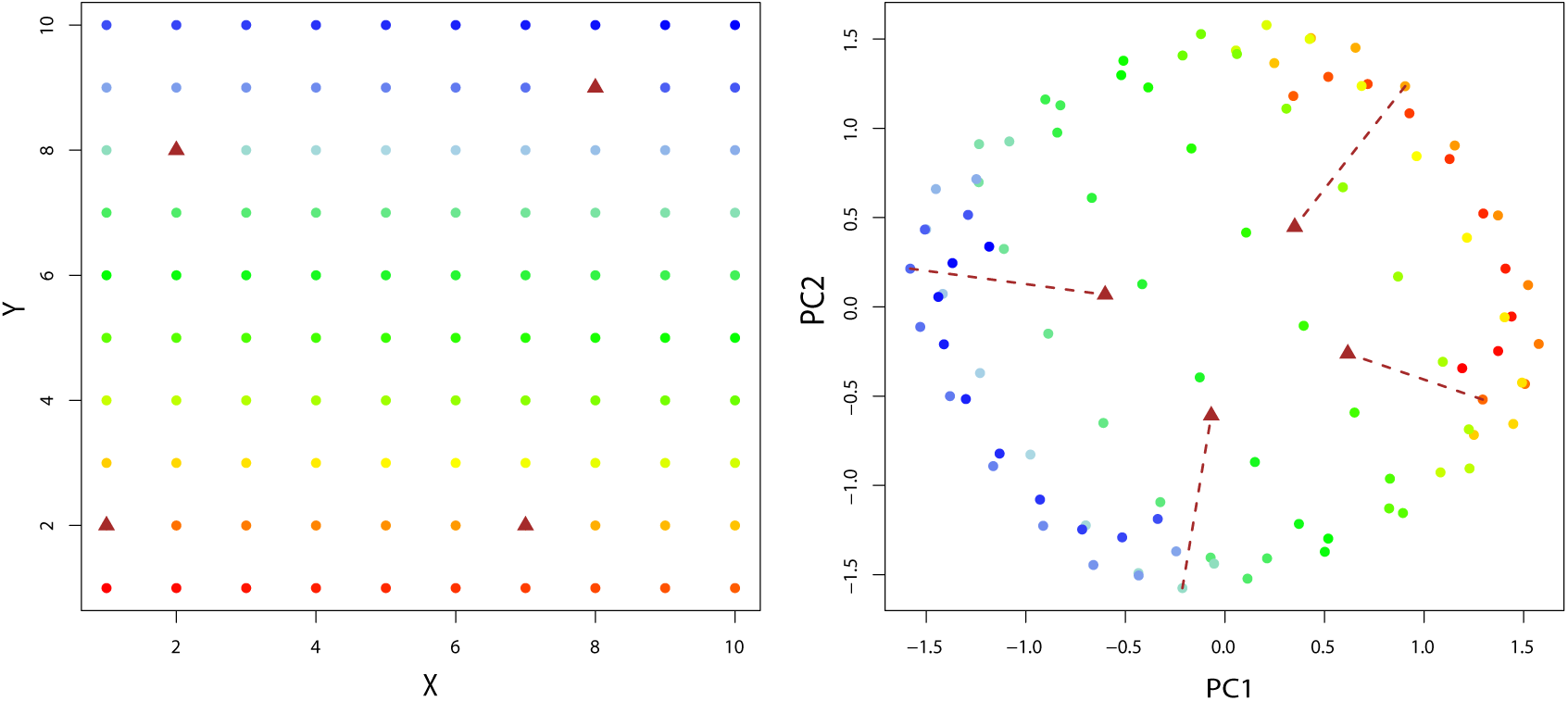
Panel A. Sampling scheme of a 10×10 grid of demes. Brown triangles represent demes where ancient individuals are sampled 1000 generations in the past. Panel B. First 2 eigenvectors of the covariance matrix between populations of Panel A. Parameters used are *m*_1_ = .01 and *m*_*∞*_ = 10^−5^. Color code is the same as in Panel A. Brown arrows start from the position of the present deme where an ancient sample is taken, and end where the ancient sample is projected on the principal components.

Using 100 demes from a 1-Dimensional simulation described above, we apply PCA to the allele frequencies at the 6000 simulated loci. To remove the edge effect, we simulate 200 demes, and consider only the 100 demes in the center. We also include allele frequencies from past generations for several demes. PC1 shows the 1-Dimensional pattern of isolation–by–distance as expected, and ancient samples are closer to 0 (Figure 6A). The distance between the scores of ancient individuals and the center of the principal component decreases as the sampling time increases. In practice, the true allele frequencies are not known, and the covariance matrix is estimated on individuals. When working with sampled individuals instead of allele frequency, the same pattern is still visible. A subsampling of 10 diploid individuals for each deme at the present time, and 1 diploid individual for each ancient deme shows the same shrinkage of PC scores for ancient individuals (Figure 6B).

**Figure 6:**
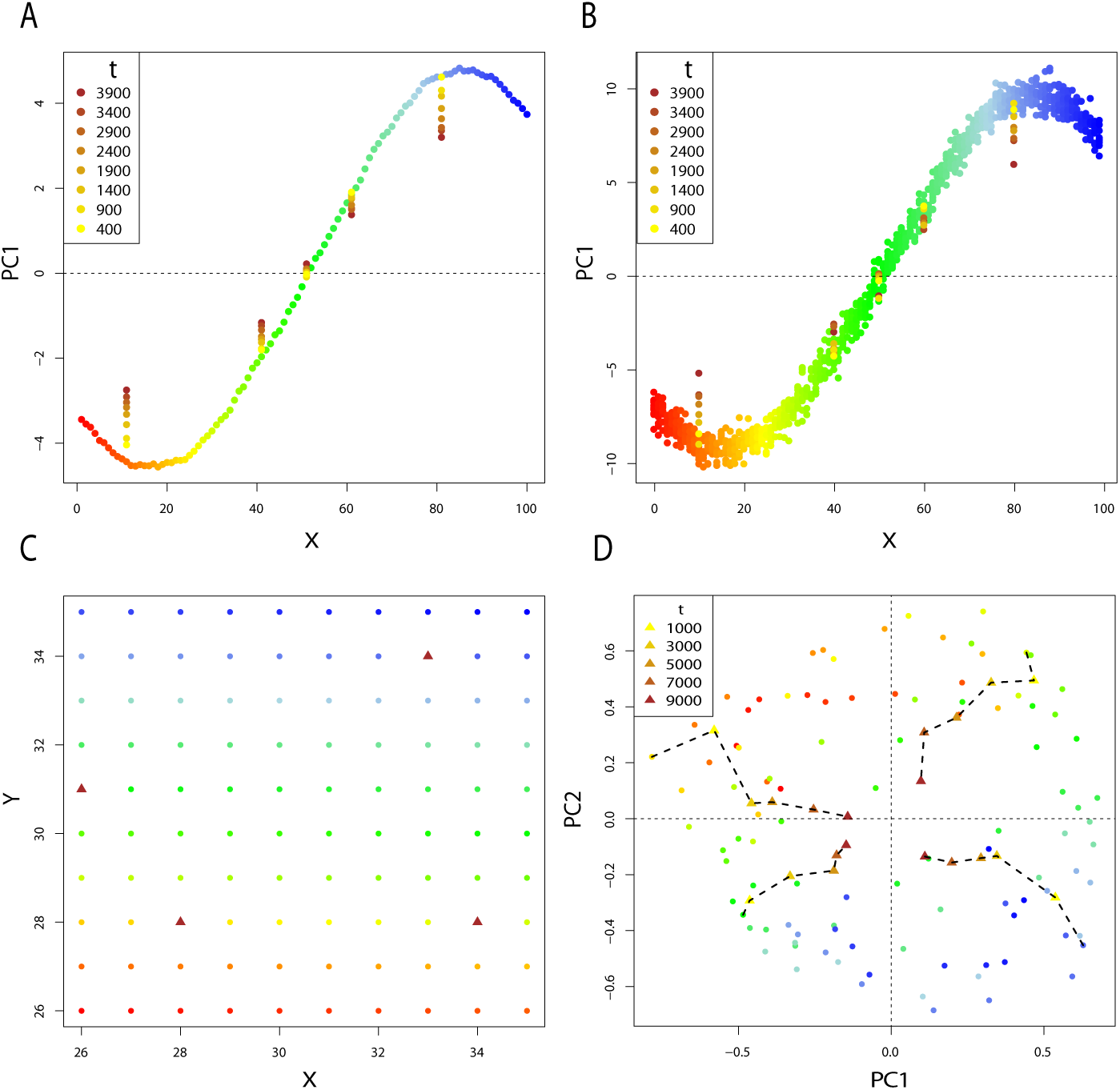
Panel A. First principal component for the 1-Dimensional simulation described above, with *m*_1_ = .01 and *m*_*∞*_ = 4.10^−5^. PCA is performed on allele frequency data from each of the 100 demes, and ancient allele frequencies are taken in 5 populations at 8 times in the past. Panel B. First principal component for the 1-Dimensional simulation described above. In each deme, 10 diploid individuals are sampled at the present time. One diploid individual is sampled in 5 demes at 8 times in the past. Panel C. Sampling scheme of a 10 × 10 grid of populations. Demes marked by a triangle are demes where ancient individuals were sampled. Panel D. plot of *P C*1 and *P C*2 for the 2-Dimensional simulation with *m*_1_ = .001 and *m*_*∞*_ = 10^−5^. Ancient samples are taken at different times in the past for 4 demes.

When applying PCA on allele frequencies from the 2-Dimensional simulations, the time effect is visible on the first two components. We study the case of a 10 × 10 grid, with no edge effects, and ancient samples taken from 4 demes at different times in the past (Figure 6C). The first and second principal components reproduce the geography of the samples, and the ancient samples are moved towards the center of the plot (Figure 6D).

This shrinkage effect of time can be understood considering the shape of the covariance function. The first and second principal components represent the 2–Dimensional Isolation–By–Distance pattern. This pattern causes the covariance matrix at time *t* = 0 to have a “block Toeplitz with Toeplitz blocks” form (Novembre and Stephens, 2008). However the pairwise covariance between present-day individuals (*t* = 0) and between ancient and present-day individuals (*t* > 0) does not have the same shape (Figure 1). Equation (10) implies that in a stepping–stone model the covariance as a function of distance flattens when comparing present and ancient individuals. As a consequence, the scores of ancient samples are moved towards the center of the principal components reproducing the local correlation pattern. Thus ancient samples can cluster with present samples at different locations, even in an equilibrium stepping–stone model.

### 5.2 One component for the time differentiation

Links between PCA and population genetics quantities, such as coalescence times and *F_ST_* have been studied (McVean, 2009; Duforet-Frebourg et al., 2015; Baran and Halperin, 2015) and show that these values can be estimated from principal components. In the 2–population case, McVean (2009) showed that the distance between individuals on the appropriate principal component is approximately a linear function of the square root of the time, ∆, until the lineages of the two individuals are in the same deme. If there are ancient and present samples, they can be considered as two groups, and ∆ is the time corresponding to the first two parts of the coalescence process between the lineages, described in the previous section. The time separating the individuals is a source of variance important enough to be reflected in the principal components (Skoglund et al., 2014). In this case, one component separates the two groups and the distance between groups is approximately proportional to 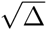. In Appendix E, we compute the expectation of ∆ if there are several present-day and one ancient individuals sampled.

We analyze the case with 50 contiguous populations sampled from a circular 1-Dimensional stepping–stone model with *n_d_* = 1000. We assume *m*_1_ = 0.1, and one deme is sampled in the past. We apply PCA by computing the eigenvectors of the individuals correlation matrix. The first principal component represents the IBD pattern between the present demes (Figure 7A). The second principal component corresponds to the differentiation between the ancient deme, and the present demes. The average distance on PC2 between the two groups (present and ancient) is an increasing function that can be approximated by a linear function of the square root of ∆ (Figure 7B).

**Figure 7:**
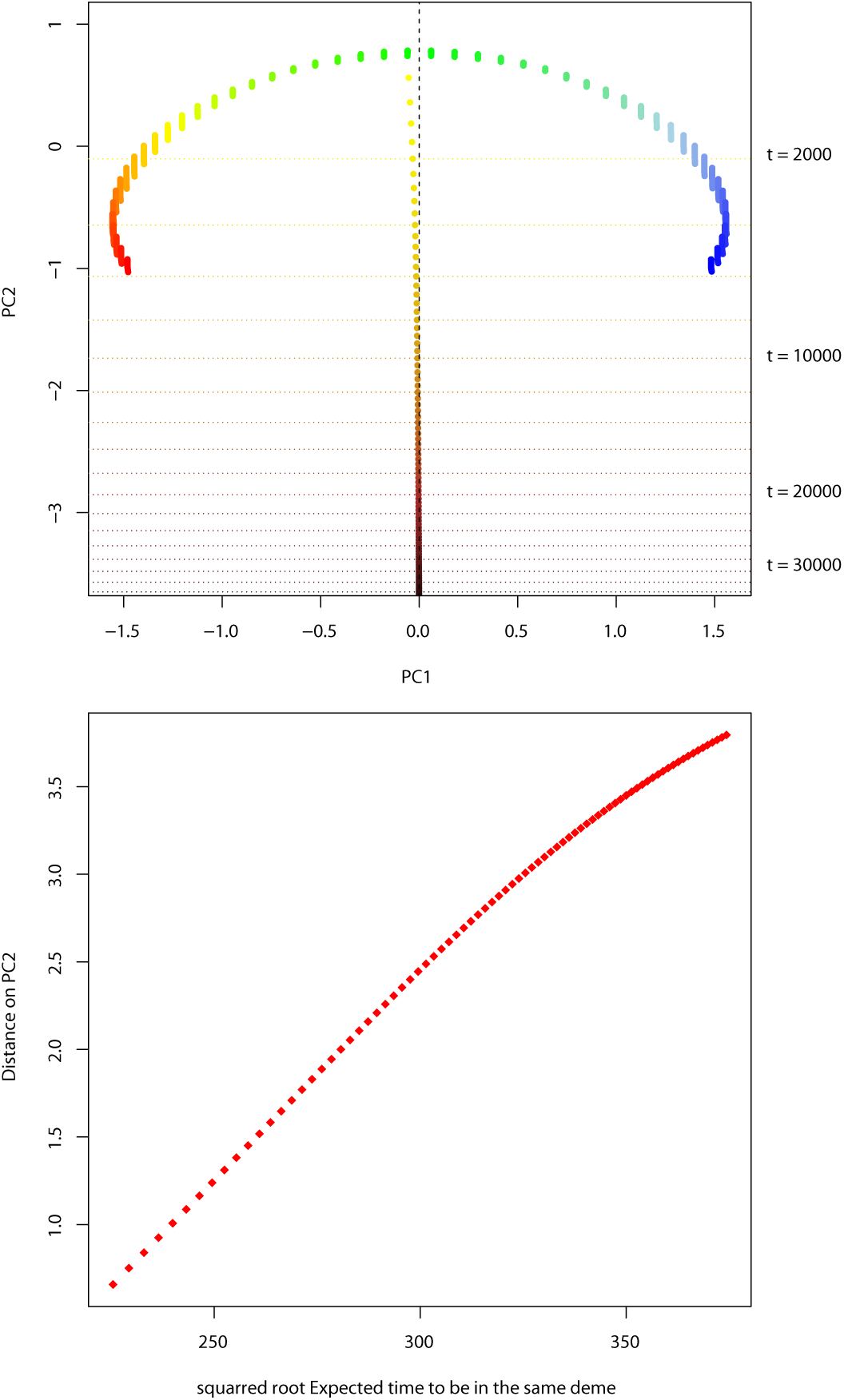
Panel A. Principal components for a 1-Dimensional stepping–stone with 50 present demes, and 1 ancient deme. The PCA is performed several times, with an ancient deme sampled at different times. Panel B. Average distance between present demes and ancient deme on PC2 as a function of 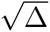

## 6 Conclusions and discussion

We have generalized the Kimura–Weiss theory of a stepping–stone model to the case where samples are taken at different times, a theory we call Isolation-by distance-and-time (IBDT). The correlation between individuals decreases as a function of both geographic distance and time. This result is accentuated in higher dimensions. When considering IBDT patterns, the edge effect applies when considering a linear model with a finite number of demes, similarly to the standard stepping–stone model. However simulations shows that in both 1 and 2 dimensions, this effect vanishes at a rate depending of the migration rate. We have also derived the expected coalescence times under the assumption of a circular, or toroidal model and low mutation rate. As the time between samples increases, the coalescence time between samples can be approximated by a linear function of time.

The connection between IBDT theory and PCA is of interest as it gives insights about what to expect from the PC plots that compare ancient and present-day samples. When considering only principal components reproducing geography, scores of the ancient samples may not cluster with the population at the same location. Such a result can occur even in the case of a population at equilibrium in a stepping–stone model, with no complex demographic history. This behavior of PCA is important to note as it could result in the inference of a non-existent ancient demographic event. The genetic differentiation created by time can be observed on another principal component. An important question that remains is in which conditions the proportion of variance explained by time is larger than the proportion of variance explained by Geography. In this unlikely event, the first principal component would not reproduce geography of the samples but rather the time line of the samples.

The limitations of PCA to investigate population structure in a spatio– temporal context highlights the need for new theoretical developments to analyze population structure when present-day and ancient samples are combined. This is especially apparent when considering the complex demographic scenarios already inferred about the history of modern humans (Pickrell and Reich, 2014). Important theoretical work has already been done to test specific hypothesis (Durand et al., 2011; Loh et al., 2013). Another way to test different past demographic events is with intensive simulation procedures, such as Approximate Bayesian Computations (Beaumont et al., 2002; Csilléry et al., 2010). In this case, theoretical developments on mechanistic models such as the stepping–stone model are important to perform simulations efficiently (Baird and Santos, 2010).

We studied the classical stepping–stone model under the assumptions of a stationary distribution of the allele frequencies in both time and space. These assumptions are not valid in all cases. The time–stationary distribution is not reached when recent events such as range expansions occurred, causing asymmetry in the site frequency spectrum (Hallatschek et al., 2007; Peter and Slatkin, 2013). Spatial non–stationarity and anisotropy can occur when migration pattern is uneven between all populations, or migration is favored in one direction (Jay et al., 2013; Duforet-Frebourg and Blum, 2014; Petkova et al., 2014). The correlation of allele frequencies is then not only a function of space and time, but also of the locations of each deme in the habitat.

A stepping–stone model is not the only model to describe spatial population structure. As an alternative to discrete models, continuous models can also be considered to study evolutionary processes (Maruyama, 1972; Barton et al., 2002, 2010). Isolation–by-Distance–and–Time can be studied in continuous framework. In the same way, results about coalescence times in a stepping–stone model can be connected to previous theory on coalescence in a continuous population (Wilkins and Wakeley, 2002).

## Acknowledgement

This work was supported by NIH grant R01-6M40282 to M. Slatkin.

## Appendix A

Using the notations in Weiss and Kimura (1965), we calculate the covariance of the allele frequencies *ρ*(*k*) between two populations that are spatially separated by *k* units of distance. This quantity is defined by

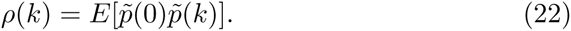

In the case where the demes are also separated by *t* units of time, we define

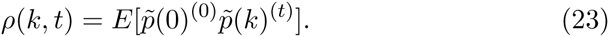

and in the particular case of *t* = 1,

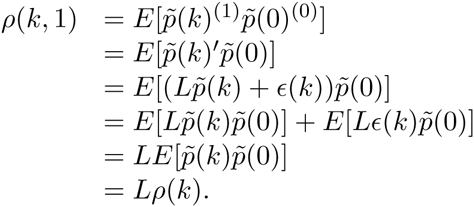

By induction, we show that for any value of *t* > 0

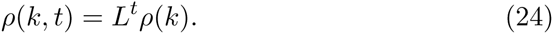

Let’s assume that for a time *t* > 0 equation (24) is true,

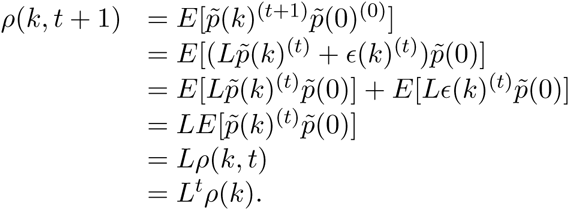

Then to obtain the correlation of allele frequencies *r*(*k, t*) between two demes, we have *ρ*(0, 0) = *ρ*(0) and

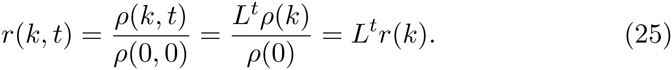

## Appendix B

We established in equation (11) that *r*(*k, t*) = *L*^*t*^*r*(*k*), and using the general expression in equation (6) we have,

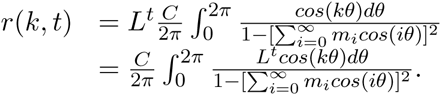

It is now demonstrated that

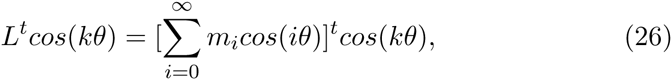

where for the convenience of the notation we denote *m*_0_ = (1 − *m*_*∞*_ − 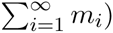. In the particular case of *t* = 1 we have

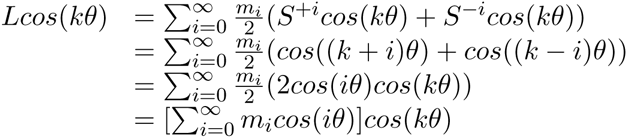

Now assuming that formula (26) holds for any value *t* > 0, we have

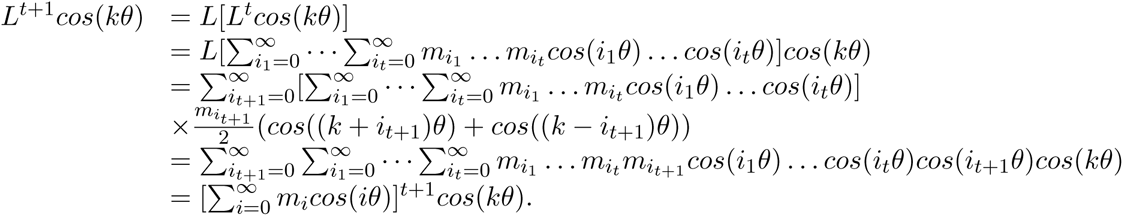

We can conclude by induction that formula (26) is true for any positive *t*. Then, using equation (26), a general formula for *r*(*k, t*) can be expressed

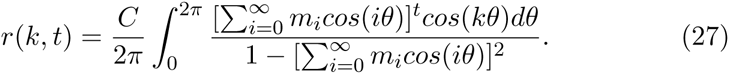

Constant C is set such that *r*(0, 0) = 1. We do not analytically investigate this constant, however details about the case *t* = 0 can be found in Weiss and Kimura (1965).

## Appendix C

Let’s assume the particular stepping-stone model: 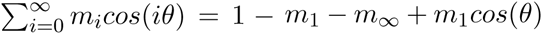. Now the correlation between 2 demes *k* steps appart and *t* generations is

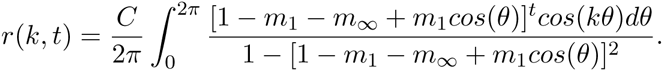

The fraction can be decomposed in two parts *r*(*k, t*) = *C*/(2*π*)(*A*_1_(*k, t*) + *A*_2_(*k, t*)) using partial fraction expansion, where

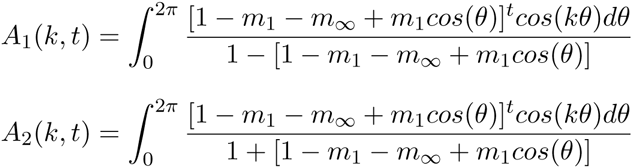

. Let *α* = (1 - *m*_1_ - *m*_*∞*_)/*m*_1_, we can expand *A*_1_ and *A*_2_,

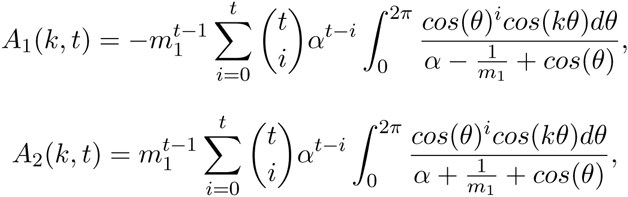

To get rid of the integral, we can use the fact that

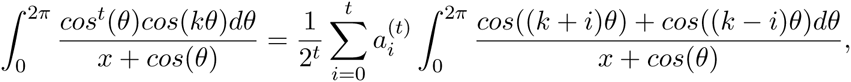

where

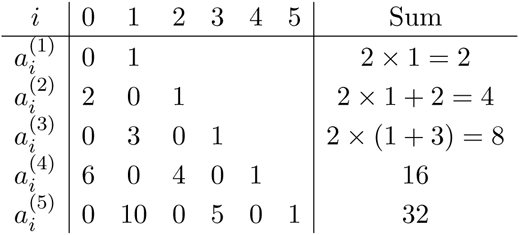

and as given in Weiss and Kimura (1965)

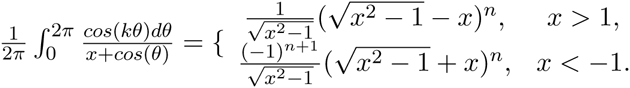

This leads us to the expressions for *A*_1_ and *A*_2_,

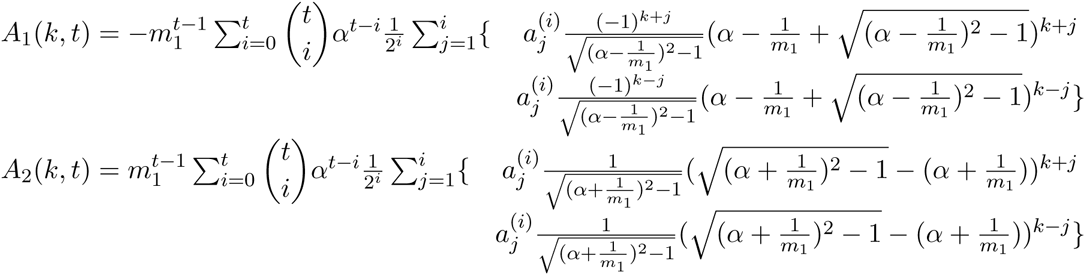

## Appendix D

The 2-Dimensional case of the analysis can be detailed by changing the operators *L* and *S*. We note the cartesian coordinates of each deme with the couple (*i*_1_, *i*_2_), and we define the operators *S*_1_ and *S*_2_ such as

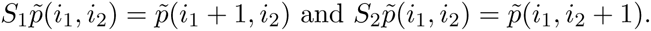

The operator *L* in two dimensions becomes

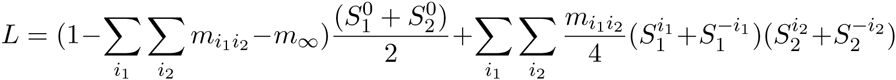

where 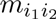 is the migration rate between demes separated by *i*_1_ and *i*_2_ steps. The correlation in 2 dimensions can be written using the spectral decomposition and for two demes we have

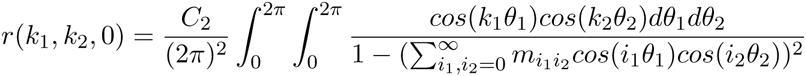

for two populations that are separated by *k*_1_ and *k*_2_ steps at the same generation. Using the same trigonometric properties as in appendix B, we have

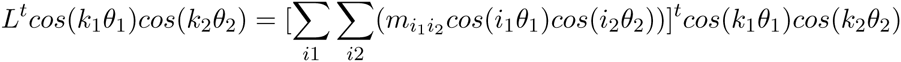

and 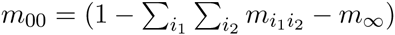. As a consequence, the correlation of allele frequencies in 2 dimensions between two populations separated by *k*_1_ and *k*_2_ steps, and *t* generations is

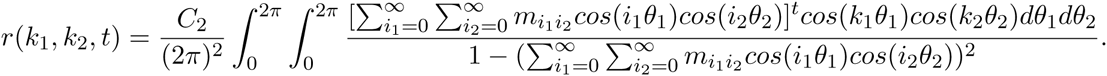

To go further, and especially investigate the 3-Dimensional case that can be relevant in practice, it is possible to extend the calculations in n-dimensional models, where two populations are separated by *t* generations and a vector of steps (*k*_1_,… *k*_n_). Redefining the operators S and L, we can show that the correlation is

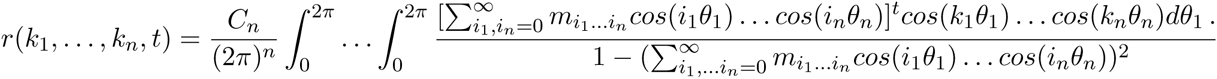

## Appendix E

We detail the case where two groups are present in the data, the present demes and the ancient deme. The quantity ∆ is the time for two genes in different groups to be in the same group. In the case where there is one ancient deme *k*_2_ and one present deme *k*_1_, using equation (19) we have

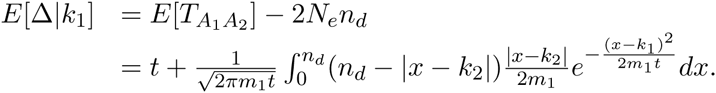

In the practical case we consider several present time demes 1… *n_p_*, and one ancient deme. The expectation of ∆ has to be conditioned by the probability that *A*_1_ is in a given present population *k*_1_.

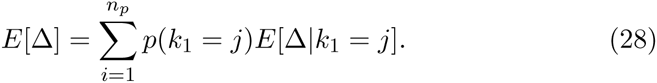

Since we consider a stepping–stone model where all the populations have the same effective population size, we have *p*(*k*_1_ = *j*) = 1*/n_p_*, *j* = 1… *n_p_*.

